# Within-host adaptive evolution is limited by genetic drift in experimental human influenza A virus infections

**DOI:** 10.64898/2026.01.07.698006

**Authors:** Lucas M. Ferreri, Nahara Vargas-Maldonado, Valerie Le Sage, Christina M. Leyson, Matthew D. Pauly, David VanInsberghe, Vedhika Raghunathan, Shamika Danzy, Hollie Macenczak, Jessica Traenkner, Flu CHIM Study Group, Andrew Catchpole, Alex Mann, Nadine G. Rouphael, Seema S. Lakdawala, Katia Koelle, Anice C. Lowen

## Abstract

Selection of advantageous mutations drives the emergence of dominant variants during seasonal influenza epidemics. However, within-host detection of such variants remains rare, limiting our understanding of how selection operates at the scale of individual hosts. In this study, we used a controlled human infection model to examine the within-host evolutionary dynamics in thirteen participants intranasally infected with a seasonal H3N2 influenza A virus. Although this clinical trial is ongoing, our work represents a pre-planned, interim, exploratory analysis. Results in this system were contrasted with those observed in a ferret model of infection. The inoculum, used in both humans and ferrets, carried standing diversity that enabled evaluation of variant trajectories during infection. Although the dynamics were variable among participants, in humans, the minor variants in the PA and NP gene segments tended to increase in frequency as infection progressed. Variant dynamics were more consistent among ferrets but showed differences from humans in the fate of the minor NP allele. Based on these observations, we fit a population genetic model to longitudinal measurements of variant frequencies. Estimates of variant selection coefficients and effective viral population sizes indicated that in humans the two minor variants had a selective advantage over the major variants, but genetic drift was strong, limiting the efficiency of selection. In ferrets, the PA minor variant also was estimated to have a selective advantage, while the NP minor variant was estimated to have a selective disadvantage. Moreover, effective viral population sizes were estimated to be considerably higher in ferrets than in humans, indicating that genetic drift was weaker in ferrets. Our analyses reveal differing selective environments acting on influenza viruses in human and ferret hosts and indicate that selection at the within-host level is weakened by genetic drift.

## Introduction

Viral variants that sweep through the human population evolve within individual hosts. Variants may carry an advantage due to immune evasion (Fitch, Leiter, Li, & Palese, 1991; Rambaut et al., 2008; Smith et al., 2004), resistance to antivirals (Ghedin et al., 2012), or increased transmission capacity (Herfst et al., 2012; Imai et al., 2012), highlighting the importance of adaptive evolution for viral spread. However, the selection of advantageous variants during the course of an acute infection has rarely been observed in individual humans. Sequencing of viral populations from naturally infected individuals has only rarely revealed mutations in known antigenic sites, for example, and has not shown these mutations to increase in frequency over the course of infection (Dinis et al., 2016).

While positive selection is apparent at the level of the host population, studies from naturally infected individuals suggest that genetic drift and purifying selection are the major evolutionary forces shaping influenza virus populations in acute infections at the within-host scale (Debbink et al., 2017; Dinis et al., 2016; J. T. McCrone et al., 2018). The strong role of genetic drift has been detected by the magnitude of variant frequency changes between longitudinally sampled viral populations. Purifying selection is apparent from the scarcity of observed nonsynonymous variants within acutely infected individuals relative to what is expected under neutral evolution. Neither of these processes readily explains the signature of positive selection shaping influenza virus dynamics at the level of the human population.

Within-host viral evolution has been examined in the context of community-acquired infections (Debbink et al., 2017; Dinis et al., 2016; Han et al., 2021; J. T. McCrone et al., 2018; Valesano et al., 2020). However, since natural infections are typically detected only after symptom onset, such studies often do not fully capture viral evolutionary dynamics (Biggerstaff et al., 2014). Additionally, seasonal influenza viruses are well adapted to the human host and are likely close to a local fitness maximum, making it difficult to identify variants that provide a fitness advantage (Sobel Leonard et al., 2016). Controlled human infections present benefits over natural infections that mitigate these limitations. The genetic composition of the initial viral population is well defined, the precise timing of infection is known, and daily longitudinal sampling enhances the precision of data collection. A previous analysis of a controlled human infection model found evidence for rapid within-host viral adaptation when variants from the inoculum stock were transferred to the participants (Sobel Leonard et al., 2016).

An important feature of influenza virus research in humans, which applies to both community cohort and experimental studies, is the complex exposure history of adult participants. Even where immune status is documented, this history is difficult to reconstruct. Animal models therefore offer a valuable complement to the study of viral dynamics in humans, since previous exposures can be eliminated or controlled. For influenza virus, ferrets are often used as a model because of their natural susceptibility to infection and onward transmission of seasonal strains (Belser et al., 2022; Bouvier & Lowen, 2010). Since host factors are often an important source of selection pressure, viral dynamics in model species may not accurately predict outcomes in humans. However, ferrets have been successfully used to evaluate within-host evolution of influenza viral populations (Moncla et al., 2016; Wilker et al., 2013).

Here, we examined the evolutionary dynamics of influenza A virus populations in experimental infections of humans and ferrets. Some of the variants tended to increase in frequency over the course of individual infections, providing initial evidence of positive selection. However, variant frequencies varied across individuals and trends in these frequencies showed differences between humans and ferrets. By fitting a population genetic model to observed variant frequencies, we found that - in both humans and ferrets - variant dynamics were best explained by the interaction between selection and genetic drift. Differences between humans and ferrets were apparent, however, in the relative fitness of viral variants and in the levels of genetic drift driving viral evolution in these two hosts. These results highlight that influenza virus populations in the upper respiratory tract can undergo positive selection during acute infection in cases when advantageous variants have high fitness and are present early in infection.

## Results

### Enrollment and experimental inoculation of healthy volunteers

Between July 2022 to October 2023, 18 adult participants were enrolled in a controlled human challenge trial at Emory University (Supplemental Figure 1)(Shetty et al., 2024; Vargas-Maldonado et al., 2025). All participants met protocol-defined eligibility criteria including a hemagglutination inhibition titer ≤1:40 against the challenge strain. On study day 1, participants were inoculated intranasally with 5.5 log_10_ median tissue culture infectious dose (TCID_50_) of influenza A/Perth/16/2009 (H3N2) (Fullen et al., 2016; Shetty et al., 2024) virus using a mucosal atomization device.

The results reported here represent a pre-planned, interim, exploratory analysis in which we evaluated the evolution of the viral population in the nasopharynx. Samples were taken daily for 7 days post inoculation from all participants. Those showing positivity by plaque assay or with Ct values equal to or less than 35 were analyzed by next generation sequencing of the viral genome. This subset of samples was derived from 13 participants, with the number of sample-days analyzed per participant ranging from 1–5. Differences in the number of samples analyzed per participant arose due to variable replication kinetics across individuals, despite the use of a controlled exposure.

### The viral inoculum carried minor variants that are prevalent in seasonal H3N2 viruses

The genetic makeup of an initial infecting population influences its evolutionary trajectory. The inoculum used in this study was derived from a stock passaged six times in embryonated eggs, which created opportunity for the virus population to diversify and diverge from the wild-type sequence. We evaluated the genetic composition of the inoculum and found four nonsynonymous mutations, two each in the PA and NP gene segments: PA I211M and I228N (frequencies of 0.36 and 0.34, respectively) and NP N101D and L136I (each with a frequency of 0.37) (Figure 1A SuppTable 1). The similar frequencies of the minor variants *within* each gene segment suggested that they are physically linked. Due to the segmented nature of the influenza virus genome, next generation sequencing is not suitable for resolving the association of mutations *between* different gene segments. Examination of clonal isolates was therefore performed. All four possible combinations of the variants at PA 211 and NP 101 were detected within isolates, at frequencies that indicate that these loci are in linkage equilibrium despite their similar observed frequencies (Figure 1B).

**Fig. 1.**
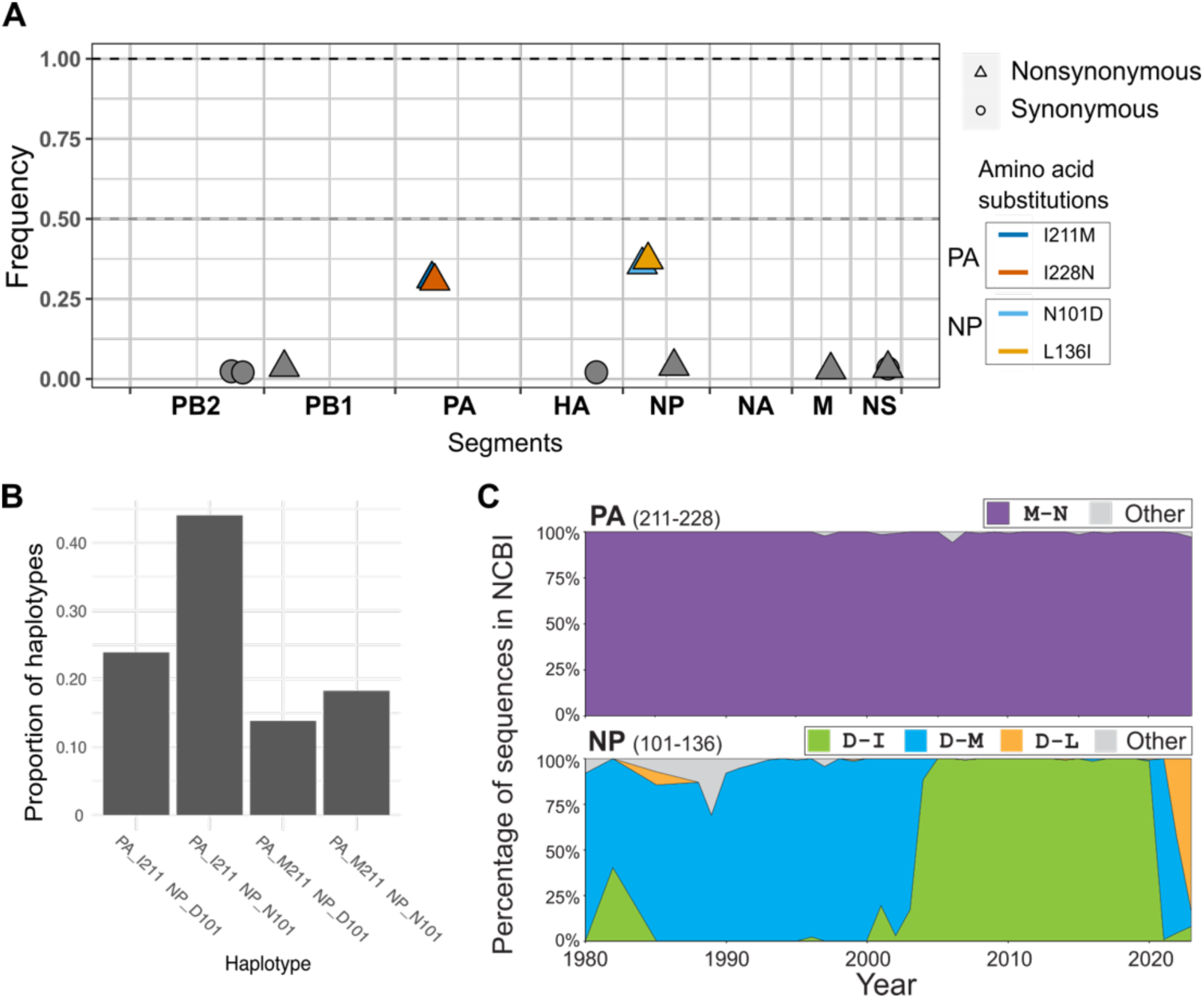
Minor variant detection in the inoculum. A. Minor variants detected in inoculum. Colored variants represent minor nonsynonymous variants within PA and NP gene segments that are the focus of this study while grey variants represent others found across the genome. B. Proportion of viral genotypes found in inoculum from 159 plaque isolates. Since the two variants within each segment are linked, here we define viral genotypes by the combination of the amino acids at sites PA 211 and NP 101 without considering PA 228 or NP 136. C. Combinations of amino acids observed in the PA gene at sites 211 and 228 (top) and in the NP gene at sites 101 and 136 in seasonal influenza H3N2 viruses over the time period 1980-2024(Ferreri & Vargas-Maldonado, 2025; Vargas-Maldonado & Ferreri, 2025).

To understand the fitness associated with these variants in nature, we examined their prevalence in the seasonal H3N2 lineage, to which the A/Perth/16/2009 (H3N2) challenge strain belongs. PA 211M, PA 228N, and NP 101D were present at frequencies close to 100% among human H3N2 strains collected between 1980 and 2024 and reported in GenBank. The amino acid at NP 136 evolved from M to I to L over this time but was predominantly I in 2009 (Figure 1C). These data indicate that the minor variants in PA and NP detected in the inoculum match the seasonal H3N2 lineage circulating in 2009. We therefore refer to these variants as wild-type.

### Minor PA and NP variants gained frequency in the human nasopharynx as infection progressed

Under positive selection, advantageous mutations increase in frequency over time. Since the wild type variants were minor variants in the challenge stock, we reasoned that they likely became minor during egg passage and may confer an advantage in participants. We therefore used next generation sequencing of viral genomes sampled from the nasal tract to evaluate the hypothesis that the wild-type PA and NP variants would increase in frequency over time (Fig. 2)(Ferreri & Vargas-Maldonado, 2025; Vargas-Maldonado & Ferreri, 2025). We observed that wild-type NP 101D-136I increased in frequency reaching high proportions in 11 out of the 13 participants (frequency range = 0.84-1). In 4 of these 11 participants, this variant was fixed (frequency >0.95). For wild-type PA 211M-228N, 6 out of the 13 participants exhibited variant frequencies exceeding 0.80 at the last time point. However, the variants in the PA and NP gene segments did not always follow similar dynamics. For instance, while the frequencies of the PA and NP variants in participants F026 and F075 exhibited similar dynamics, the frequencies of these variants in participants F031 and F048 exhibited discordant dynamics, with one wildtype allele nearing or reaching fixation, while the other remained as a minor variant (Fig. 2). Overall, these data suggest that wild-type variants in both gene segments likely confer an advantage in human infections, although their within-host dynamics vary between individuals.

**Fig. 2.**
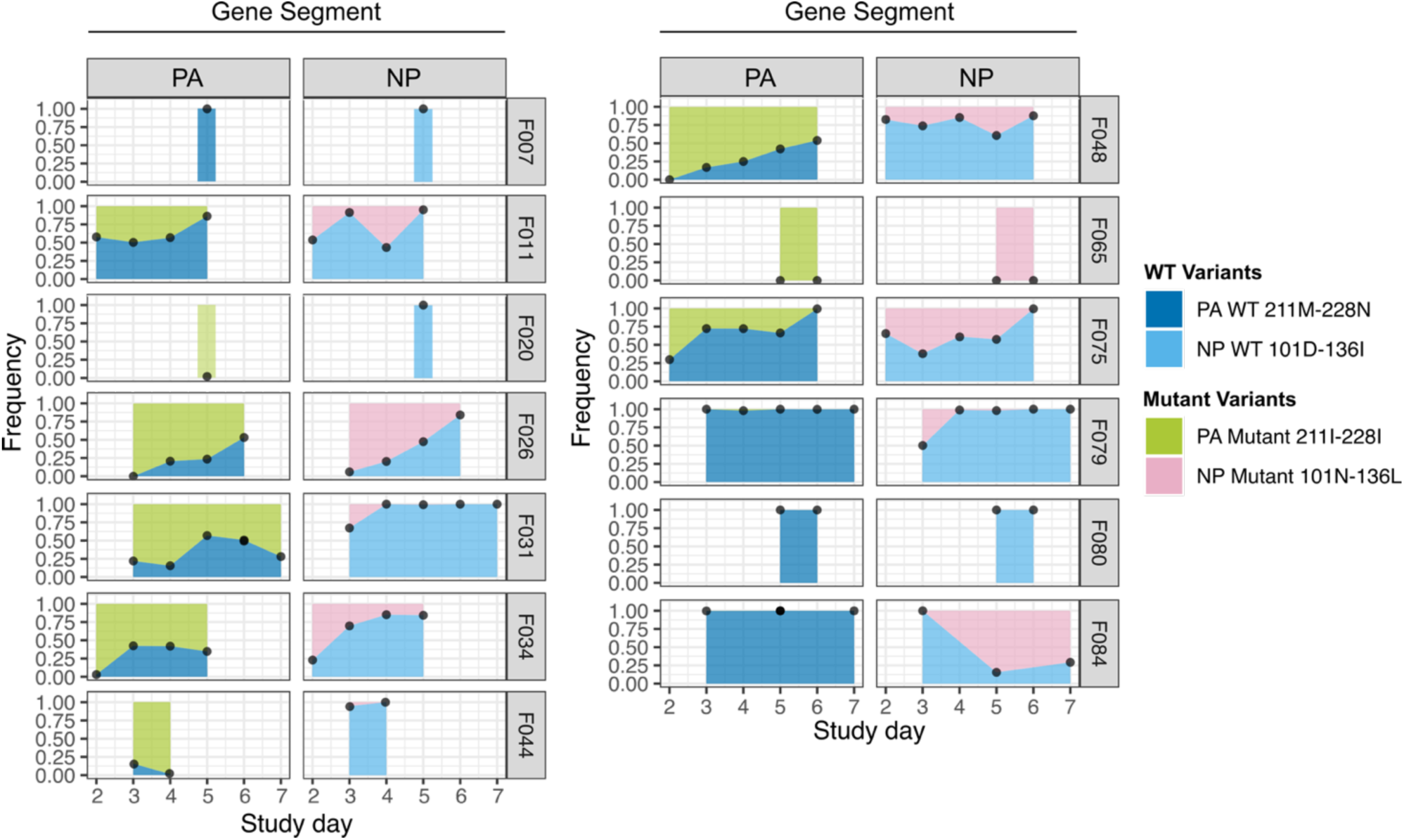
Evolution of PA and NP variants in the nasopharynx of infected humans. PA and NP variant frequencies are shown for each participant, with the participant identifier indicated at the right of each panel. Inoculation occurred on study day 1. Frequencies show the mean of the two variants within each segment (SuppTable 2, which themselves were calculated by averaging across technical replicates. Viral loads in the nasopharyngeal swabs that were processed for viral sequencing are shown in SuppFig. 2.

### Infection and transmission in ferrets show that PA and NP variants evolve in a host-dependent manner

To further evaluate the relative fitness of the viral variants present in our inoculum stock and whether these can influence transmission, we analyzed the evolutionary dynamics of these variants both within hosts and between hosts in a ferret model (Fig. 3)(Ferreri & Vargas-Maldonado, 2025; Vargas-Maldonado & Ferreri, 2025). Naïve ferrets were inoculated intranasally with the human challenge inoculum. At 24 h post-inoculation, these infected animals were placed next to additional naïve ferrets to initiate exposure. Exposure lasted two days, with the animals separated by a perforated barrier during this time. Nasal washes collected longitudinally from both inoculated and exposed ferrets were processed to evaluate viral population dynamics within and between hosts.

**Fig. 3.**
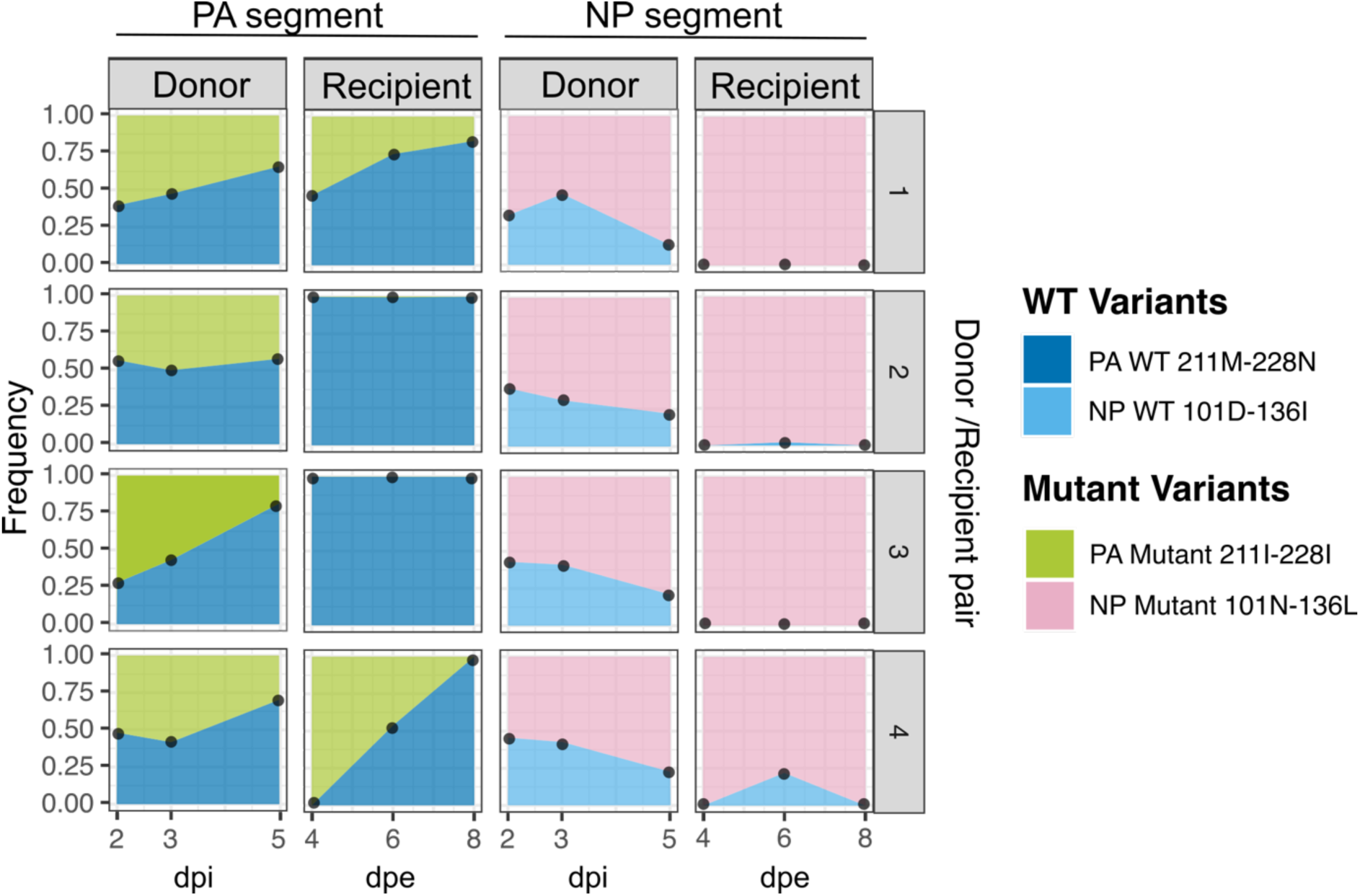
Evolution of PA and NP variants in the nasal tract of infected ferrets. PA and NP variant frequencies are shown for donor and recipient transmission pairs. Transmission pair ID numbers are shown at the right of each plot. dpi = days post infection; dpe = days post exposure. Inoculation occurred at 0 dpi and recipients were exposed to donors for 48 h beginning at 1 dpi (0 dpe). Frequencies show the mean of the two variants within each segment (SuppTable 3. There was no averaging of frequencies for a given variant because only one replicate was performed. Viral loads of nasal wash samples that were processed for viral sequencing are shown in SuppFig. 3.

The dynamics of PA and NP variants were largely consistent across ferrets. In directly inoculated animals, wild-type PA 211M-228N increased in frequency as the infection progressed in three out of four ferrets and maintained a frequency of approximately 50% throughout the infection in the fourth ferret. After transmission, PA 211M-228N variants were fixed from the first time point onward in two recipient ferrets and showed an upward trajectory in the other two recipients. These within- and between-host variant frequency changes indicate that the wild-type variant PA 211M-228N likely had a fitness advantage in ferrets, consistent with our findings in humans.

The variants in NP, in contrast, evolved differently in ferrets compared to humans. In donor ferrets, wild-type NP 101D-136I consistently decreased in frequency between 2 and 5 days post-inoculation. After transmission, NP 101D-136I was observed only at low levels in two recipient animals, while remaining below the limit of detection in the other two. These data suggest that, in contrast to humans, NP 101D-136I is likely deleterious in ferrets.

### Genetic drift interacts with selection in shaping the fate of variants within human participants

To more quantitatively evaluate the strength and direction of selection of the PA and NP variants in the human challenge study, we estimated the selection coefficients (*σ*) of the wild-type PA 211M-228N and NP 101D-136I variants using their observed frequencies over time. Because the observed trajectories varied across individuals (Fig. 2), we reasoned that genetic drift may also impact frequency dynamics. We therefore jointly estimated selection coefficients and the effective viral population size, *N*_e_. *N*_e_ quantifies the extent of genetic drift in a population, with smaller values indicating higher levels of genetic drift. Our joint estimation of *N*_e_ and *α* for PA 211M-228N indicates that this wild-type variant appears to have a selective advantage over the major variant PA 211I-228I (Fig. 4A), with a maximum likelihood estimate for the selection coefficient being *σ* = 0.070, corresponding to 7.3% higher fitness. The 95% confidence interval for the selection coefficient ([-0.015, 0.190]) does cross zero, however. We further find support for a very small effective population size, with an estimate of *N*_e_ = 19 (95% CI = [13, 33]) viral particles. The 95% confidence region shown in Fig. 4A further indicates that our model, incorporating both selection and genetic drift, is likely to be statistically preferred over an analogous model that incorporates only selection (*N*_e_ = ∞) or only genetic drift (*σ* = 0). Indeed, when we compare across these three models using the Akaike Information Criterion corrected for small sample size (AIC_c_), we find that the selection plus drift model is preferred over both the selection-only model and the genetic drift-only model (SuppTable 4), albeit only slightly.

**Fig. 4.**
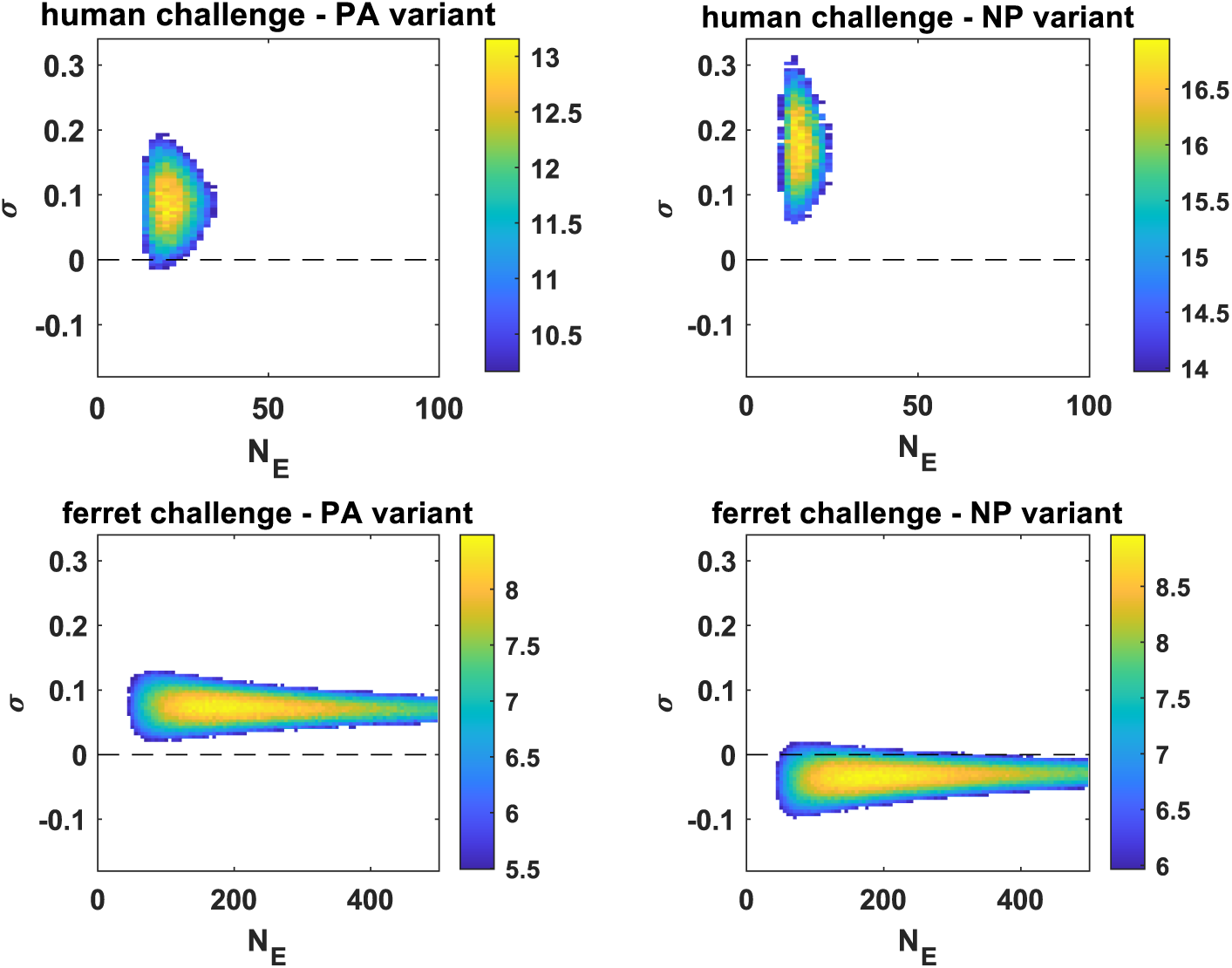
Joint estimates of variant selection coefficients and effective viral population sizes. (A) Estimates based on frequency changes of PA 211M-228N observed in the 13 participants. (B) Estimates based on frequency changes of NP 101D-136I observed in the 13 participants. Frequency changes in PA 211M-228N and in NP 101D-136I in these participants are shown in Fig. 2. (C) Estimates based on frequency changes of PA 211M-228N observed in directly inoculated (donor) ferrets. (D) Estimates based on frequency changes of NP 101D-136I observed in directly inoculated (donor) ferrets. Frequency changes in PA 211M-228N and in NP 101D-136I in the donor ferrets are shown in Fig. 3. In (A)-(D), only log-likelihood values of parameter combinations that fall within the 95% confidence region are shown. Color bars to the right of the panels show the range of calculated log-likelihood values. Supplementary Fig. 4 and 5 show model-generated reconstructions of variant frequencies for the experimentally challenged humans and ferrets, respectively. Supplementary Fig 6 and 7 show the log-likelihood profiles for the genetic drift-only model and for the selection-only model.

Our joint estimation of *N*_e_ and *α* for NP 101D-136I in humans indicates that this wild-type variant also has a selective advantage over the major variants NP 101N-136L (Fig. 4B), with an estimate for the selection coefficient being *σ* = 0.190 (95% CI = [0.055, 0.310]), corresponding to 21.0% higher fitness. We again find support for a very small effective population size, with an estimate of *N*_e_ = 15 (95% CI = [9, 23]) viral particles. When we compare the selection-and-drift model against a selection-only and a genetic drift-only model for this NP variant, we this time find that the selection plus drift model is statistically strongly preferred over the selection-only model and the genetic drift-only model (SuppTable 4).

In conclusion, we estimate that wild-type PA 211M-228N has a moderate selective advantage over the mutant PA 211I-228I and that wild-type NP 101D-136I has a relatively strong selective advantage over the mutant NP 101N-136L in human infections. Furthermore, we find that genetic drift plays a large role in shaping the frequencies of these variants in the controlled human challenge study, with estimates of *N*_e_ being very small. The strong role of genetic drift that we infer explains why, despite the selective advantage of these wild-type variants, there is considerable variation observed across individuals.

### Selection and, to a lesser extent, genetic drift shapes the fate of variants within ferrets

We next applied the same population genetic model to infer the strength of selection and genetic drift acting on the PA and NP variants in the ferret challenge study. Our joint estimation of *N*_e_ and *α* for PA 211M-228N indicates that this wild-type variant again has a selective advantage over the major variant PA 211I-228I, this time in ferrets (Fig. 4C), with the estimate for the selection coefficient being *σ* = 0.065 (95% CI = [0.020, 0.125]), corresponding to 6.7% higher fitness. We find support for a moderately sized effective population size, with an estimate of *N*_e_ = 168 (95% CI = [43, 1290]) viral particles. When we compare across selection-plus-drift, selection-only, and drift-only models using AICc, we find that the selection-plus-drift model is again statistically preferred over the selection-only model and the genetic drift-only model (SuppTable 4).

In contrast, our joint estimation of *N*_e_ and *α* for NP 101D-136I indicates that this wild-type variant appears to have a selective disadvantage over the major variant NP 101N-136L (Fig. 4D), with the maximum likelihood selection coefficient estimated to be *σ* = -0.035. This corresponds to a fitness of NP 101D-136I that is 96.6% of that of NP 101N-136L. The 95% confidence interval for the selection coefficient ([-0.100, 0.015]) does cross zero, however. We again find support for a moderately sized effective population size, with an estimate of *N*_e_ = 158 (95% CI = [43, 2730]) viral particles. When we compare the selection-and-drift model against a selection-only and a genetic drift-only model using AICc, we again find that the selection-plus-drift model is preferred over the selection-only model and the genetic drift-only model (SuppTable 4), albeit only slightly over the genetic drift-only model.

In conclusion, in ferrets, we estimate that PA 211M-228N has a moderate selective advantage over PA 211I-228I and that NP 101D-136I has a weak selective disadvantage compared to NP 101N-136L. These results indicate that a variant that is favorable in one host may be unfavorable in another host and that genetic drift appears to be weaker in ferrets relative to humans in experimental infections. These findings highlight the presence of host-specific differences in the direction of selection and the efficiency with which selection can act within experimentally infected hosts.

## Discussion

Leveraging the standing diversity of the inoculum, we evaluated the within-host evolutionary dynamics of influenza A virus populations in humans and ferrets challenged with the same virus stock. Our results reveal evidence of selection in both systems, with the variant PA 211M-228N tending to increase in frequency over time in both hosts and the variant NP 101D-136I tending to increase in humans but decrease in frequency over time in ferrets. Variant dynamics were, however, variable among individuals in both the human and the ferret challenge studies, indicating a degree of evolutionary stochasticity. In line with these observations, quantitative analyses indicated that variant dynamics in both humans and ferrets could be best explained through a combination of selection and genetic drift. Nevertheless, differences between human and ferret hosts were apparent, with selection favoring wild-type NP 101D-136I in humans but mutant NP 101N-136L in ferrets and genetic drift being more prominent in humans than in ferrets.

Our observation of positive selection acting during the course of acute infection differs from many prior reports on the within-host evolution of influenza viruses. Previous studies examining natural influenza virus infections in humans have shown that genetic drift and purifying selection are dominant forces (Debbink et al., 2017; Dinis et al., 2016; J. T. McCrone et al., 2018; Sobel Leonard et al., 2016; Valesano et al., 2020). In contrast, signatures of positive selection acting on seasonal influenza viruses are readily detected at the scale of the host population (Smith et al., 2004; Thompson et al., 2024).

Examination of the differences between natural, acute infections and the experimental infections evaluated here suggests some likely reasons for the rarity of positive selection within naturally infected hosts. First, while our experimental inoculations used a large initial population, natural influenza virus infections are thought to be established by relatively small founding populations (J. T. McCrone et al., 2018; T. Shi, Harris, Martin, & Koelle, 2023). This increases the extent of genetic drift at early timepoints following infection in naturally infected individuals. Second, time limitations – due to the acute nature of a typical influenza virus infection - are likely to have a larger impact when variants arise *de novo* than when they are delivered with the inoculum. The importance of the timescale of infection for the likelihood of viral adaptation is supported by the documentation of positive selection during prolonged influenza virus infections in immunocompromised individuals (Rogers et al., 2015; Xue et al., 2017). Third, since seasonal influenza viruses are well adapted to the human host, the distribution of mutational fitness effects is likely to be heavily skewed towards negative effects (Visher, Whitefield, McCrone, Fitzsimmons, & Lauring, 2016), such that mutations arising during natural infection are unlikely to confer an advantage. Thus, the NP and PA alleles present in our challenge stock offered an unusual opportunity to monitor the efficiency of selection acting on advantageous variants. In summary, the genetic variation in the inoculum used in this study, along with a large founding viral population size at the start of the experimental infections, likely enabled the occurrence of within-host selection. Of note, Sobel Leonard and colleagues have also reported adaptive evolution during acute influenza virus infection in experimentally inoculated humans (Sobel Leonard et al., 2016). Unlike in our study, these variants were nonsynonymous variants on the hemagglutinin gene segment. This study referred to these evolutionary dynamics as purifying selection on the (mutant) inoculum stock variants, rather than positive selection on the wild-type variants, but these two forces are equivalent, differing only in the allele whose dynamics are being followed. Although observed in artificial systems, both of these studies demonstrate the potential for adaptive influenza virus evolution to occur during the course of acute infection under favorable circumstances.

While selection could be detected, the dynamics of the PA and NP variants in our data sets also show evidence of genetic drift. These stochastic effects were quantified through estimation of the effective population size *N*_e_, revealing that only a few dozen members of the viral populations in humans and only a few hundred members of these viral populations in ferrets contribute progeny to the next generation during each within-host infection cycle. The *N*_e_ values that we estimate for humans are consistent with the small effective viral population sizes estimated in two studies analyzing allele frequency dynamics from natural human infections (John T. McCrone, Woods, Monto, Martin, & Lauring, 2020; Y. T. Shi, Martin, Weissman, & Koelle, 2025), but lower than recent estimates from another study that estimated *N*_e_ for natural influenza infections of 284 ± 60 for a seasonal influenza A H3N2 virus and 176 ± 41 for the 2009 pandemic influenza A H1N1 virus (Bendall et al., 2024). These latter differences likely relate to methods of variant calling applied. We are not aware of prior estimates of *N*_e_ for influenza A virus infections in ferrets.

When considered in the context of the large viral populations that form within the first 2-3 days of infection, effective population sizes on the order of a few dozen to a few hundred indicate that the progeny of only a small minority of viruses are successful *in vivo*. This feature of infection likely results in part from the high heterogeneity of viral burst size seen at a single-cell level (Bacsik et al., 2023; Heldt, Kupke, Dorl, Reichl, & Frensing, 2015; Jacobs et al., 2019). Indeed, most cells infected with a single influenza A virus fail to replicate the full viral genome (Brooke et al., 2013; Jacobs et al., 2019), resulting in cellular infections that yield no infectious viral progeny. Heterogeneity at the level of infected tissues is also likely to contribute to low *N*_e_: due to limits on the number of target cells in a local area, dispersal to new sites within the respiratory tract can create uneven opportunities for viruses to expand. Within-host dispersal to the lower respiratory tract in particular shows strong genetic bottlenecks, suggesting that relatively few members of the source population seed infection in the lungs (Amato et al., 2022; Ferreri et al., 2025). Together with the effects of superinfection exclusion (Sims et al., 2023), spatial constraints within hosts reduce the likelihood that fitter variants will be propagated and have the opportunity to outcompete less fit variants (Gifford, de Visser, & Wahl, 2013). Finally, host responses to infection that suppress the production of viral progeny are likely to contribute to low *N*_e_. As with infection, these responses show strong heterogeneity at single-cell and tissue levels (Russell, Elshina, Kowalsky, Te Velthuis, & Bloom, 2019; Sun et al., 2020), such that the immune suppression of viral populations is likely to be uneven and give rise to stochastic losses of within-host viral lineages. In sum, we propose that single-cell, spatial and immune heterogeneity each contribute to low viral effective population sizes.

Our comparison of influenza virus infection in humans and ferrets provides insight into the evolutionary processes in each system. Directional trends in variant frequencies, indicative of selection, showed less variability across individual ferrets than across individual humans, an observation which is quantitatively reflected in the higher *N*_e_ estimates for ferrets. Additionally, the selection coefficients of certain variants indicated qualitatively different selective pressures are active in humans and ferrets. The potential drivers of these differences in viral evolutionary dynamics are many. Although both humans and ferrets are highly susceptible to seasonal influenza virus infection (Belser, Katz, & Tumpey, 2011; Carrat et al., 2008), divergence in the host factors which support or impede viral replication are likely to yield differing selective pressures and viral variant dynamics. In addition to species differences, disparity in immune memory may be important: while the ferrets were immunologically naive, the human participants likely had multiple prior exposures to influenza. The resultant differences in the nature and kinetics of adaptive immune responses could precipitate differences both in the strength of genetic drift and the nature of selective pressures. Additional factors could relate to the method of inoculation. In humans, the virus was delivered using an atomizer whereas in ferrets, the virus was administered via nasal instillation. These differences may potentially yield distinct distributions of the viral populations in the two hosts and therefore differential access to target cells.

Certain limitations of our research are important to consider. These include the examination of viral evolutionary dynamics during experimental, rather than natural, infection. While some of the resultant differences enable the detection of evolutionary forces – as discussed above – others may confound translation of our findings to natural systems. In addition, the inherent limitations of next generation sequencing for the analysis of low titer viral samples and the detection of minor variants hinder our ability to decipher frequency dynamics early and late in infection and at times when alternative alleles are rare (J. T. McCrone & Lauring, 2016). Finally, while sequencing replicates, longitudinal samples and the examination of multiple individuals offer means of validating the trends observed, a lack of replicate nasopharyngeal samples prevents analysis of the contributions of sampling noise to the variant dynamics observed.

In summary, our analyses show that positive selection is active during experimental seasonal influenza virus infection of the upper respiratory tract in humans and ferrets, but the interplay with genetic drift reduces its effectiveness. Since the epidemiology of seasonal influenza is directly linked to its evolution, these findings have implications for infection burdens (Pybus & Rambaut, 2009). Adaptation of viral populations in the upper respiratory tract may promote the transmission of beneficial variants, increasing the number of individuals who become infected during an influenza season. However, the strength of genetic drift—during transmission or dispersal within the host—is likely to reduce the probability that adaptive mutations are successfully transferred. The balance between these two evolutionary forces shapes how influenza virus populations evolve and, in turn, how they spread through human populations.

## Materials and Methods

### Clincial trial information

Studies in a Controlled Human Infection Model (ClinicalTrials.gov identifier NCT05332899) were conducted at Emory University Hospital from July 2022 to October 2023 under Emory IRB protocol STUDY00000083. Before participating, all individuals provided informed consent and successfully completed an assessment to confirm their understanding of the study procedures and risks.

The study included six cohorts comprising a total of eighteen participants, each of whom received an intranasal inoculation of the seasonal influenza A/Perth/16/2009 (H3N2) virus. The virus, manufactured by Meridian Life Sciences (Memphis, TN) on behalf of hVIVO (FDA IND #19579), was administered at a concentration of 5.5 log₁₀ TCID₅₀/mL. A total volume of 0.5 mL was delivered to each nostril using an Intranasal Mucosal Atomization Device (MAD Nasal) while participants remained in a supine position.

Participants were screened for hemagglutinin inhibition (HAI) antibodies against the challenge strain up to 60 days before inoculation, as described previously (Shetty et al., 2024). Those with HAI titers of ≤40 underwent additional pre-enrollment evaluations, including a medical history review, physical examination, laboratory tests, electrocardiogram (ECG), a multiplex upper respiratory pathogen panel (BioFire, Salt Lake City, UT), and chest radiograph imaging.

On study day 0, participants were admitted to the hospital and placed under an eight-day quarantine.

### Biosafety and biocontainment

Because influenza A virus is transmissible, all challenge activities were conducted with biocontainment procedures designed to mitigate risk to participants, staff, and the community. Participants were admitted to the hospital and managed in an inpatient quarantine setting during the post-inoculation period (eight-day quarantine in this study). The challenge virus was a previously circulating influenza A/H3N2 strain and was handled and administered only in the quarantine unit. Storage and preparation followed controlled procedures (single-use vials stored at −70°C to −80°C; preparation by research pharmacy personnel). At admission, participants underwent respiratory pathogen screening using a multiplex panel, and viral shedding was monitored daily during quarantine by PCR. Discharge timing and post-discharge guidance were structured to reduce onward transmission risk, including education to avoid contact with individuals at high risk for influenza complications after release and provision of oseltamivir in participants without two consecutive negative PCR tests by discharge or with severe influenza. Only healthy participants were eligible. Healthcare personnel used personal protective equipment as part of unit procedures and received seasonal influenza vaccines.

### Nasopharyngeal sampling methodology

Nasopharyngeal (NP) sampling was performed on the day of admission and then daily from study days 2 to 8. Unless the initial swab was fully saturated with fluid, a single swab was used to collect specimens from both nostrils. In cases where a deviated septum or blockage made sampling from one nostril difficult, the same swab was used to collect the specimen from the other nostril. After collection, the swab was placed in a transport tube containing 1 mL of viral transport media (BD Universal Viral Transport Collection Kit, Cat. No. 220526, Fisher Scientific). Samples were stored at 2°C–8°C before processing for quantitative PCR (qPCR) and subsequently preserved at −80°C for infectious virus analysis.

### Animal ethics statement

The ferret experiment was conducted at the University of Pittsburgh in compliance with the guidelines of the Institutional Animal Care and Use Committee under approved protocol 22061230. Ferrets were sedated using isoflurane for all nasal washes and survival blood draw procedures, as directed by approved methods.

### Ferret screening

Immunologically naïve ferrets were screened for antibodies against influenza A and B viruses prior to purchase using sera provided by Triple F Farms (Sayre, PA) (Le Sage et al., 2025). All ferrets tested negative for previous influenza exposure.

### Transmission study design

Four five-month-old male ferrets were inoculated intranasally with the CHIM inoculum at the same dose as was used in humans, 5.5 log₁₀ TCID₅₀/mL, with 0.5 mL administered per nostril. For the 2-day exposure, each of the four infected donor ferrets was housed on one side of a one-inch-thick perforated divider and a single recipient ferret was placed on the other side, creating four 1:1 transmission pairs. Exposure was initiated at 24 hours post-donor infection. Constant directional airflow traveled from the donor to the recipient. Once the exposure complete, the recipients were individually housed for the remainder of the study. Body weight and clinical signs were recorded upon collection of nasal wash samples on the indicated days.

### Characterization of viral PA-NP genotypes present in the viral inoculum

Residual A/Perth/16/2009 (H3N2) virus from the inoculum stock was used to infect a confluent monolayer of MDCK cells grown in 10 cm dishes. The inoculum was diluted such that each plate contained 16-50 plaque-forming units (pfu) per dish. After a 1 hour incubation, agar-containing overlay was placed on the inoculated cells. Dishes were incubated for 48 hours to allow plaques to form. Agar plugs containing well-isolated plaques, i.e. those that were separated by a minimum of 0.5 cm from other plaques, were placed in 160 uL of sterile phosphate-buffered saline (PBS). This sample was then subjected to RNA extraction using the ZR-96 viral RNA kit (Zymo, USA) per the manufacturer’s protocol. Reverse transcription (RT) was performed to generate cDNAs using Maxima RT (Thermofisher, USA) and UnivF(A) universal forward primer for influenza (5’-GCG CGC AGC AAA AGC AGG-3’). Each cDNA generated from plaque-derived RNA was diluted 1:4 in nuclease-free water and used as template for NP- or PA-specific PCR using Precision Melt Supermix (Bio-Rad, USA) and CFX384 Touch real-time PCR detection system (Bio-Rad). High-resolution melt analysis was then used to distinguish the single nucleotide changes G637T (corresponding to M211I) in the PA segment or G324A (corresponding to D101N) in the NP segment. The following primer pairs were used for PA and NP segments respectively: (1) Perth_PA_602_F: 5’-AGTCCGAAAGAGGCGAAGAA -’3, Perth_PA_691_R: 5’-AGTTCGGTGGGAGACTTTGG-3’; (2) Perth_NP_281_F: 5’-TGGAAGAACACCCCAGCG-3’, Perth_NP_407_R: 5’-CGCCAGATTCGCCTTATTTCT-3’. Precision Melt Analysis software (Bio-Rad) was then used to analyze melting curves. The genotype of each plaque was assigned according to melt curves obtained from control plasmids carrying either the parental or the mutagenized sequence. Generation of control templates matching Perth09 PA and NP alternative alleles was performed using site-directed mutagenesis on pHW plasmids containing the Perth09 PA or NP segments. The mutations made correspond to the amino acid changes M211I (G637T) on the PA segment or D101N (G324A) on the NP segment. Sanger sequencing was used to confirm the presence of the desired change. These templates were then used as controls in high resolution melt analysis-based genotyping of plaque isolates derived from the inoculum.

### Whole Genome Amplification

Viral RNA was extracted as previously described. Reverse transcription and amplification were performed using 2.5 μL of RNA in a 25 μL MS-RT-PCR reaction (2), utilizing the Superscript III High-Fidelity RT-PCR Kit (ThermoFisher). The primers used and their concentrations were: Uni12/Inf-1 (5’-GGGGGGAGCAAAAGCAGG-3’) at 0.06 μM, Uni12/Inf-3 (5’-GGGGGGAGCGAAAGCAGG-3’) at 0.14 μM, and Uni13/Inf-1 (5’-CGGGTTATTAGTAGAAACAAGG-3’) at 0.2 μM.

Thermal cycling conditions were as follows: 55°C for 2 minutes, 42°C for 60 minutes, and 94°C for 2 minutes; followed by 5 cycles of 94°C for 30 seconds, 44°C for 30 seconds, and 68°C for 3.5 minutes; then 26 cycles of 94°C for 30 seconds, 55°C for 30 seconds, and 68°C for 3.5 minutes; with a final extension at 68°C for 10 minutes.

Amplicons were sequenced at either the Emory National Primate Research Center Genomics Core or the Emory Integrated Genomics Core. Sequencing libraries were prepared using the Nextera XT DNA Library Preparation Kit (Illumina) according to the manufacturer’s instructions. Libraries were multiplexed and sequenced on an Illumina NovaSeq 6000 platform using a paired-end 100 nucleotide run configuration.

### Variant Analysis

Non-consensus variant analysis was performed using a custom pipeline based on LoFreq (Wilm et al., 2012). Sequencing adapters were removed with Cutadapt, and reads were aligned to the reference genome using BWA-MEM (Li & Durbin, 2009). Subsequent data processing included mate information correction with Samtools and probabilistic realignment using LoFreq Viterbi to address mapping errors. Reads were then sorted with Samtools, and indel quality scores were added using LoFreq indelqual. Base and indel alignment qualities were incorporated using LoFreq alnqual.

Variant calling was conducted with LoFreq, producing a VCF file. Custom scripts were employed to filter and reformat the VCF, extracting allele frequencies and coverage statistics. We used a variant calling threshold of 1% and sequencing depth of at least 400X. To determine synonymous and nonsynonymous changes, SNPdat (Doran & Creevey, 2013) was used.

### Estimation of selection coefficients and the strength of genetic drift

We estimated effective viral population size (*N*_e_) and the selection coefficient (*σ*) using longitudinal measurements of variant frequencies. We estimated *N*_e_ and *α* separately for each of the two variants (PA 211M-228N and NP 101D-136I) and for each host (human and ferret). Estimation of *N*_e_ and *α* used a basic bootstrap particle filter (Chopin, 2020). The underlying state-space model was a Wright-Fisher model with selection, parameterized with a given *N*_e_ and a selection coefficient *σ* of the wild-type variants relative to the mutant variants. We set the fitness of mutant variants PA 211I-228I and NP 101N-136L both to one. Fitness values of the wild-type variants PA 211M-228N and NP 101D-136I are given by exp(*σ*), such that *σ* values below 0 correspond to deleterious variants and *σ* values above 0 correspond to beneficial variants. When estimating *N*_e_ and *α* using frequency changes of PA 211M-228N, for each particle in the bootstrap particle filter, we set each initial virion (of the *N*_e_) to be 211M-228N with probability 0.3505, corresponding to the frequency of this variant in the inoculum stock. Similarly, when estimating *N*_e_ and *α* using frequency changes of NP 101D-136I, for each particle, we set each initial virion to be 101D-136I with probability 0.3727, corresponding to the frequency of this variant in the inoculum stock. As such, we assume that there was a loose bottleneck in the establishment of infection upon challenge. Variant frequencies were simulated from one viral generation to the next using the Wright-Fisher model with selection assuming a viral generation time of 6 hours. Particle weights were calculated at every timepoint at which variant frequencies were observed. The measurement model to calculate the particle weights was a normal distribution with a standard deviation of 0.05, truncated between the frequencies of 0 and 1 and renormalized.

Particles were resampled using a multinomial resampling scheme. Likelihoods of parameter combination (*N*_e_, *α*) were determined by first calculating the arithmetic means of the particle weights at each observation timepoint and then taking the product of these means across all observation timepoints, as is routinely done in a bootstrap particle filter. We used 1000 particles in the bootstrap particle filter and simulated the bootstrap filter 10 times for each parameter combination. Log-likelihood values shown in Figure 4 are means of log-likelihood values across these 10 independent bootstrap filters. Variant frequencies were reconstructed for each bootstrap filter by sampling a particle at random at the end of each simulation. By independently estimating *N*_e_ and *α* for each wild-type variant, we implicitly assume that reassortment is common between the PA and NP gene segments, such that the frequency changes of these variants are due to selection only on their own gene segments. To ensure that our estimates were robust, we excluded human samples from our analysis if their Ct values were above 30 or if their viral titers fell below 10^1^ PFU/mL. This included day 7 of participant F031, day 2 of participant F048, day 2 of participant F075, days 3, 5, 6, and 7 of participant F079, days 5 and 6 of participant F080, and day 7 of participant F084. Estimation of *N*_e_ under the drift-only model was performed analogously to that for the selection-and-drift model, except that the selection coefficient *σ* was set to 0 and log-likelihoods were calculated across a range of *N*_e_ values.

Estimation of *σ* under the selection-only model was also performed analogously to that for the selection-and-drift model, except that the effective population size *N*_e_ was set to 100,000 and only one particle (instead of 1000) and only one bootstrap filter (instead of 10) was used. An *N*_e_ of 100,000 was sufficiently large to yield evolutionary dynamics that appeared deterministic (as would be in the absence of genetic drift). Because of these deterministic dynamics, only a single particle was needed and only a single bootstrap filter.

## Supporting information

SuppTable 1

SuppTable 2

SuppTable 3

## Data availability

Next-generation sequencing data can be found in BioProjects PRJNA1367170 and PRJNA1196946 of the Sequence Read Archive (SRA). Code for variant analysis is available on Github: https://github.com/genferreri/Initial_LoFreq. Code for estimating the effective viral population size and the selection coefficient under all three models (drift, selection, and selection-and-drift) is available on GitHub: https://github.com/koellelab/within_host_IAV_drift_and_selection Data on viral titers, variant frequencies, and modeling results presented in this manuscript are available on FigShare at DOI: 10.6084/m9.figshare.30801677.

## Acknowledgements

This study was supported in part by Flu Lab and the National Institute of Allergy and Infectious Diseases (NIAID) through U01 AI144673 and the Centers of Excellence for Influenza Research and Response (CEIRR) contract no. 75N93021C00017. We acknowledge the valuable contributions of the members of the Emory University Flu CHIM Study Group: Meredith J. Shephard, Michelle N. Vu, A.J. Campbell, Kayla Brizuela, Ralph Tanios, Veronica Smith, Cecilia Zhang, Kareem Bechnak, Sarah Bechnak, and Jianguo Xu.

## Author Contributions

Conceptualization: LF, NVM, NGR, SSL, KK, ACL

Clinical trial design: NGR, SSL

Clinical study implementation: NGR, SSL, MDP, HM, JT, Flu CHIM

Study Group Essential reagents: AM, AC

Data collection: NVM, VR, SD, CML, VLS

Data analysis: LMF, NVM, DV, KK, VLS

Preparation of initial manuscript draft: LF, KK, NVM, ACL

Review and editing of manuscript: All authors

Funding acquisition: ACL, SSL, NGR, KK

## Author Declaration of Interest

NGR receives funding from Merck, Sanofi, Pfizer, Vaccine Company, Immorna, and consulting fees from Krog &Partners. Merck, CSK is a consultant for Ferring Pharmaceuticals. LCM serves as a consultant for MITRE corporation. None of these funders or consulting agencies were involved in the research described or influenced the studies.

---

**Supplemental figure 1.**
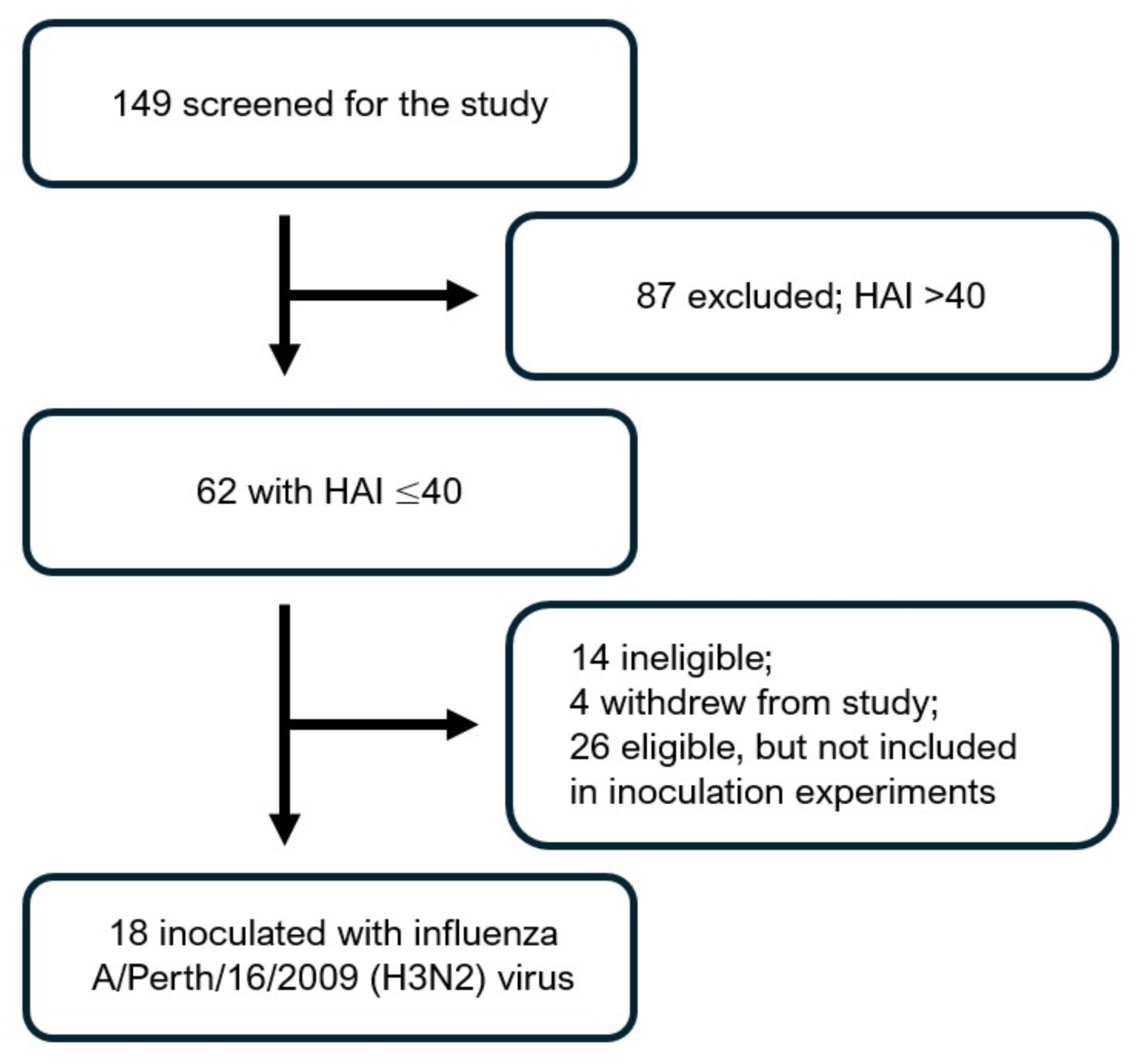
Consort diagram summarizing the recruitment and enrollment in the study.

**Supplementary figure 2.**
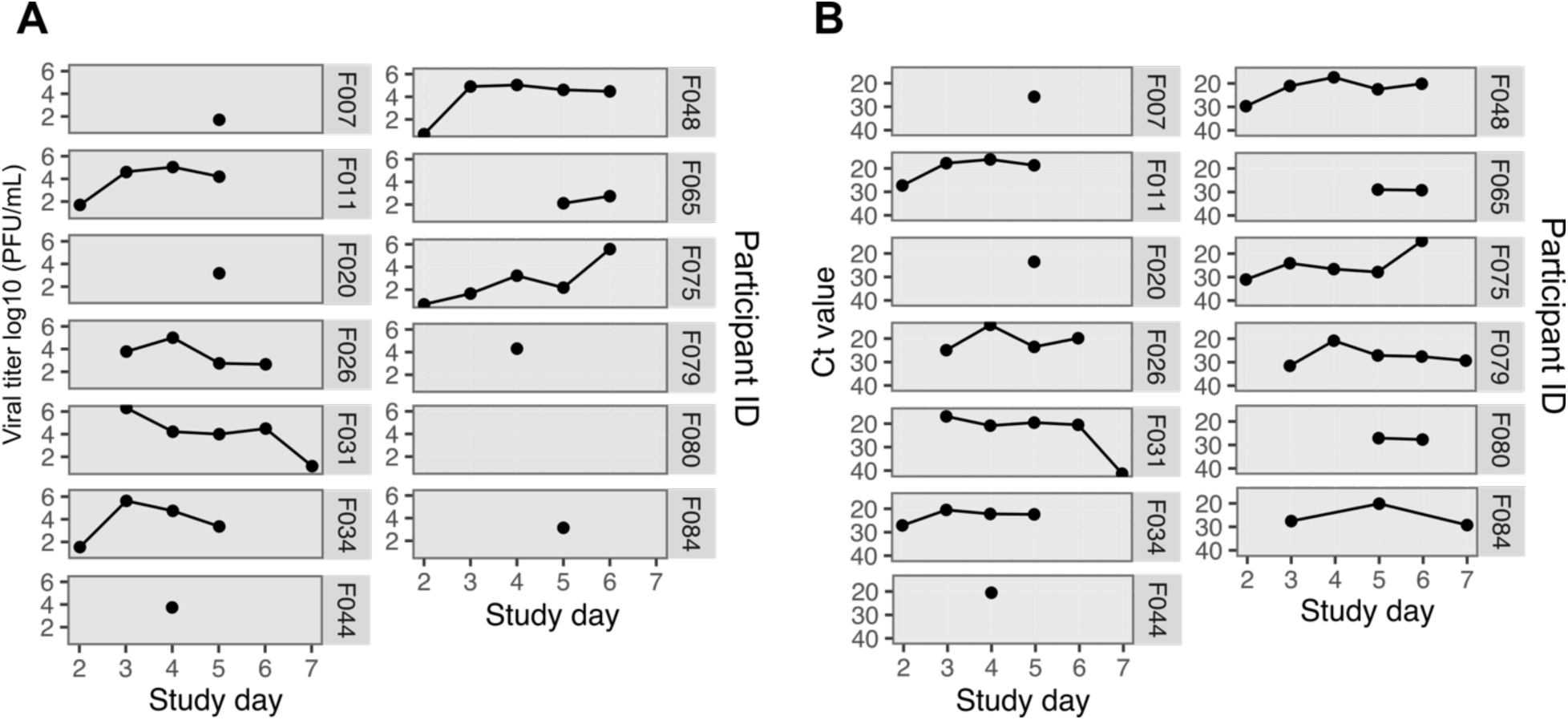
Viral loads and RNA detection in swabs from the nasopharynx. Each panel represents a single participant. Participant ID is denoted at the right of each panel. A. Viral titers as determined by plaque assay. B. Quantification of viral RNA by qRT-PCR.

**Supplementary figure 3.**
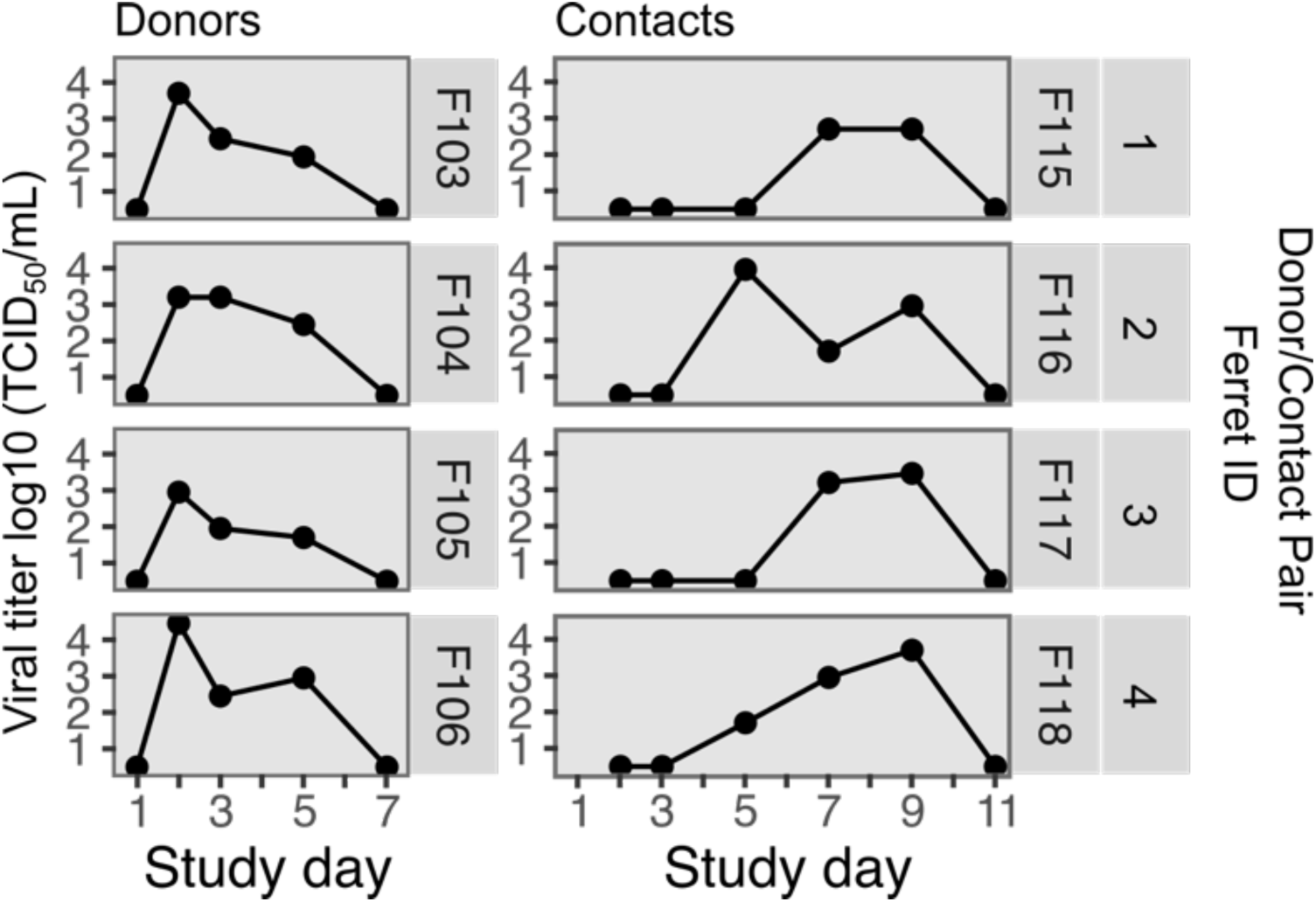
Viral loads quantified from nasal washes ofinoculated donor ferrets and their contacts. Each panel represents a single ferret. Ferret ID along with donor/contact pair are denoted at the right of each panel.

**Supplementary figure 4.**
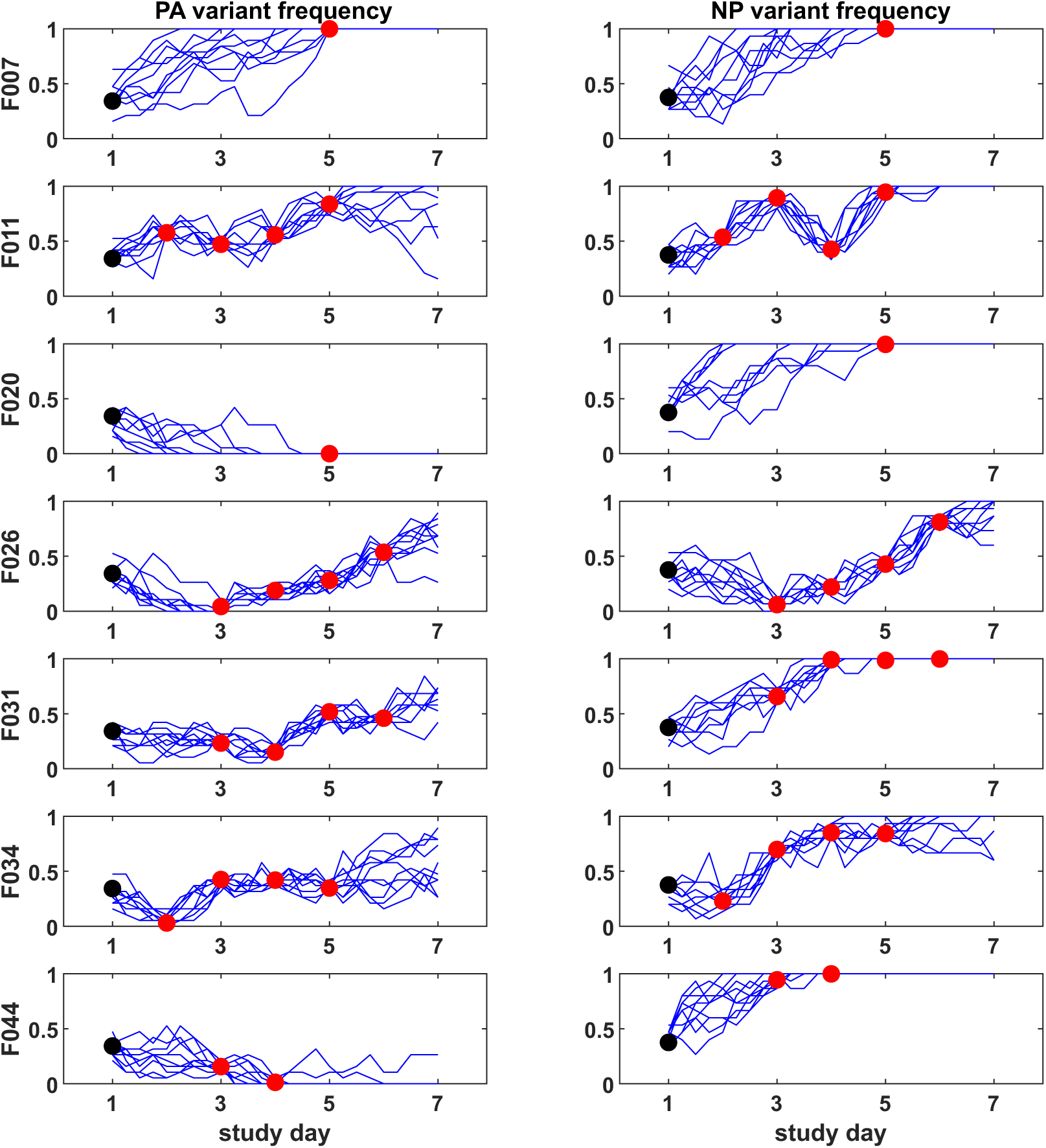
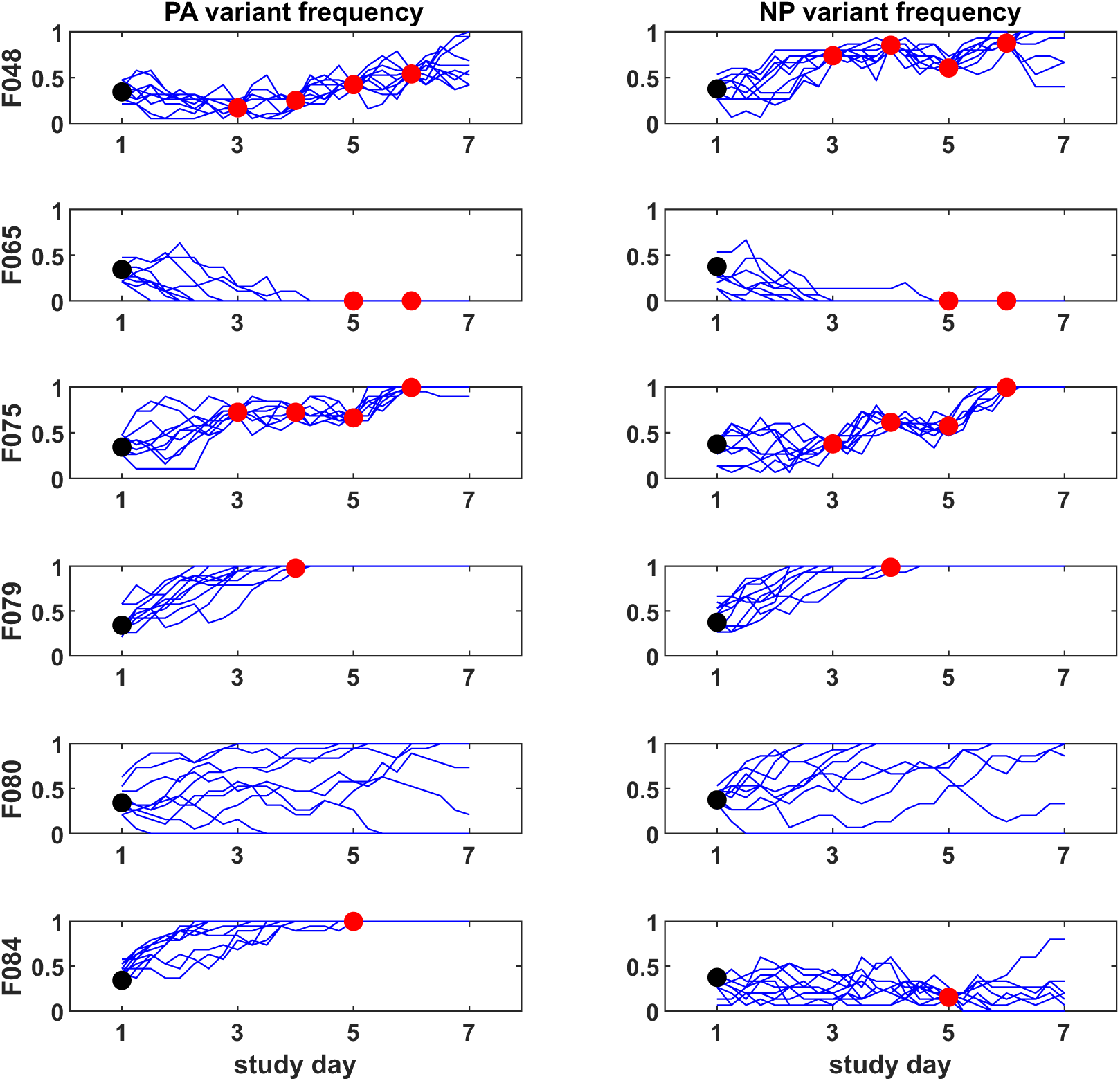
Model-generated reconstructions of variant frequencies in humans. Participants and variants are organized as in Figure 2. The black point shows the frequency of variants present in the inoculum and the red points show the frequencies observed within the host. Only timepoint samples included in the population genetic analyses are plotted. Blue lines show the frequencies reconstructed by the selection and drift model.

**Supplementary figure 5.**
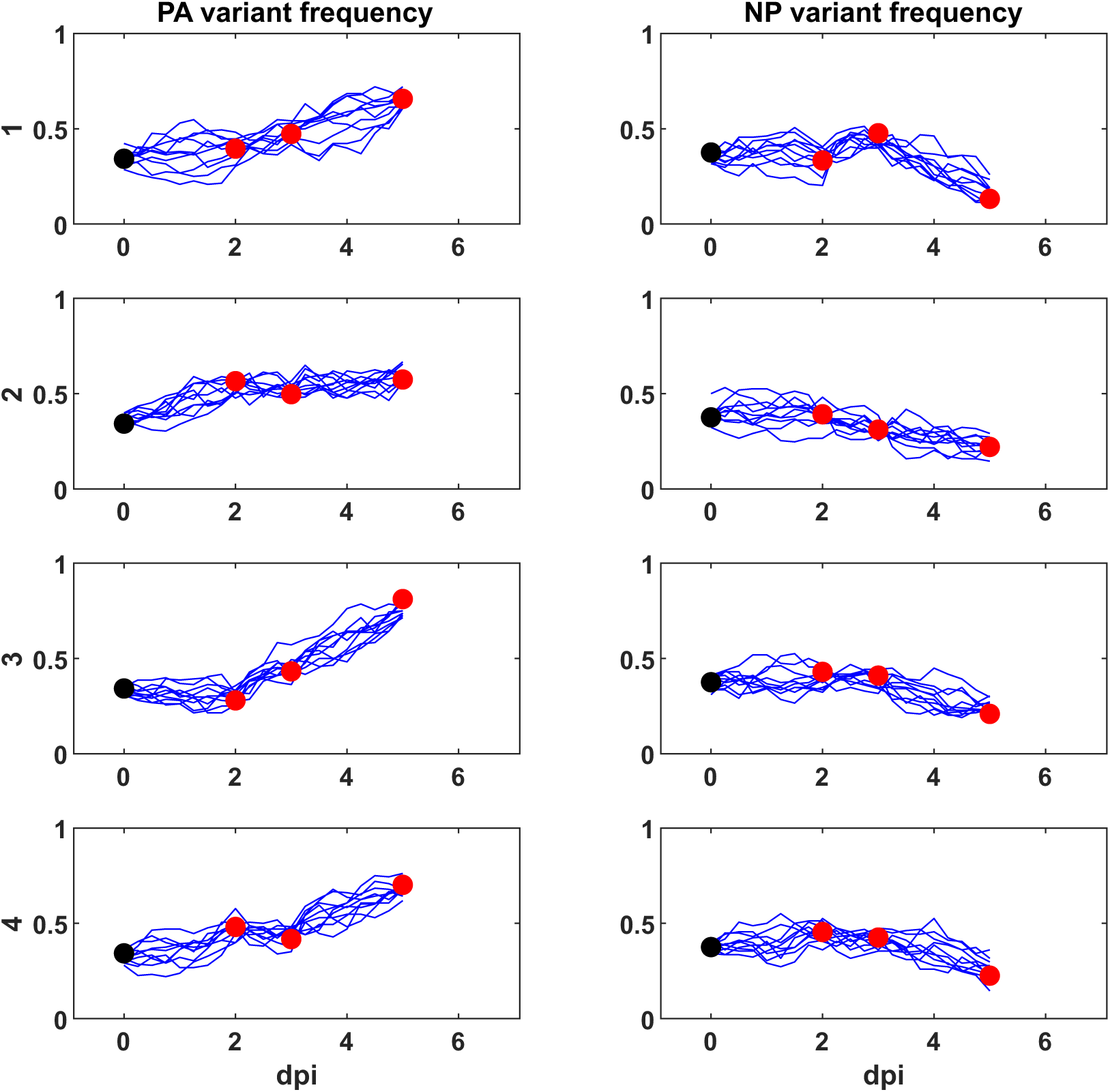
Model-generated reconstructions of variant frequencies in donor ferrets. Donor ferrets are organized as in Figure 3.Points represent observed variant frequencies, as shown in Figure 3.The black point shows the frequency of variants in the inoculum, and the red points show frequencies observed within the donor ferret. Blue lines show the frequencies reconstructed by the selection-and-drift model.

**Supplementary figure 6.**
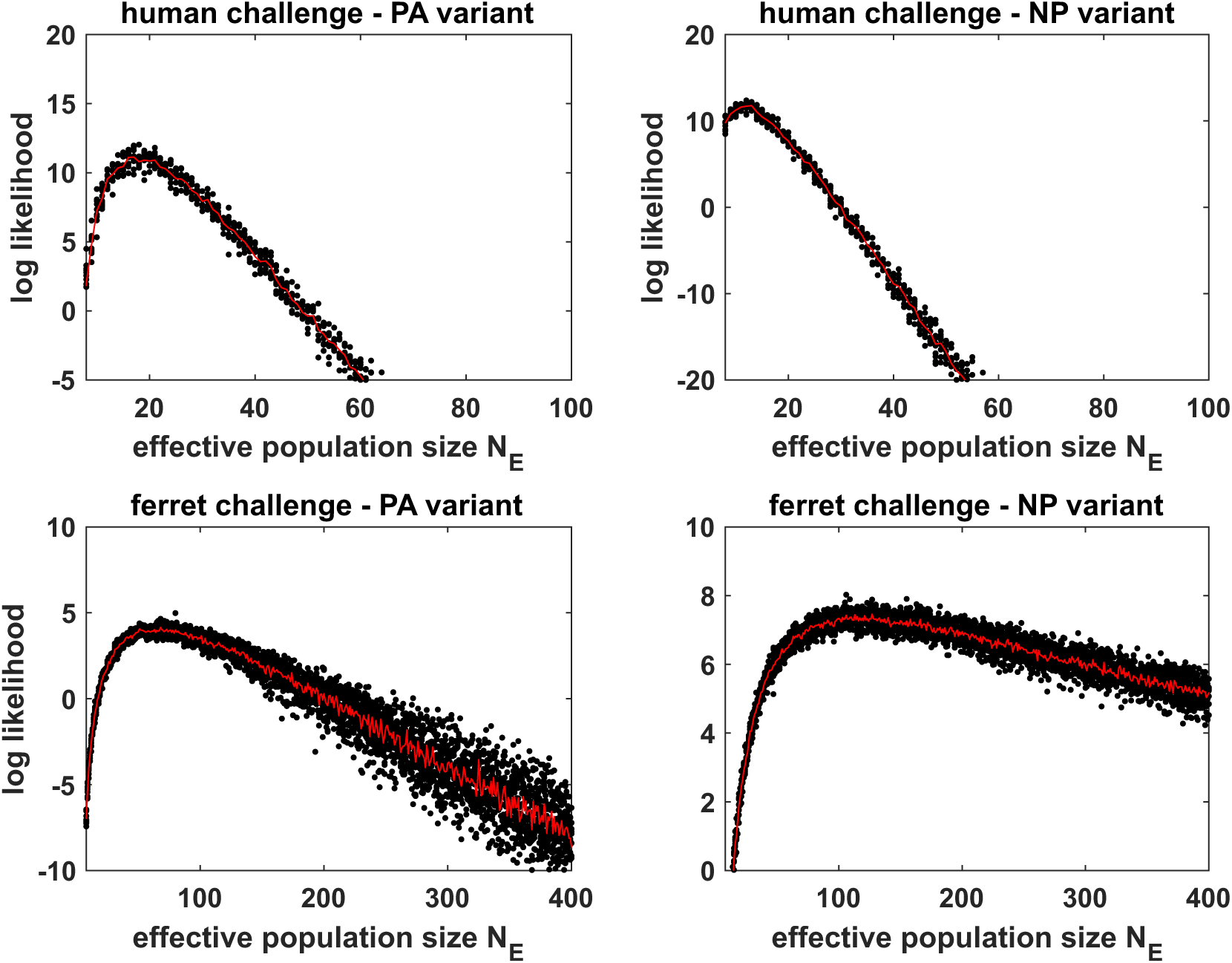
Log-likelihood profiles for effective viral population sizes, by host and variant for the genetic drift-only model. (A) Estimates based on frequency changes of PA 211M-228N observed in the 13 participants. (B) Estimates based on frequency changes of NP 101D-136I observed in the 13 participants. (C) Estimates based on frequency changes of PA 211M-228N observed in directly inoculated (donor) ferrets. (D) Estimates based on frequency changes of NP 101D-136I observed in directly inoculated (donor) ferrets.

**Supplementary figure 7.**
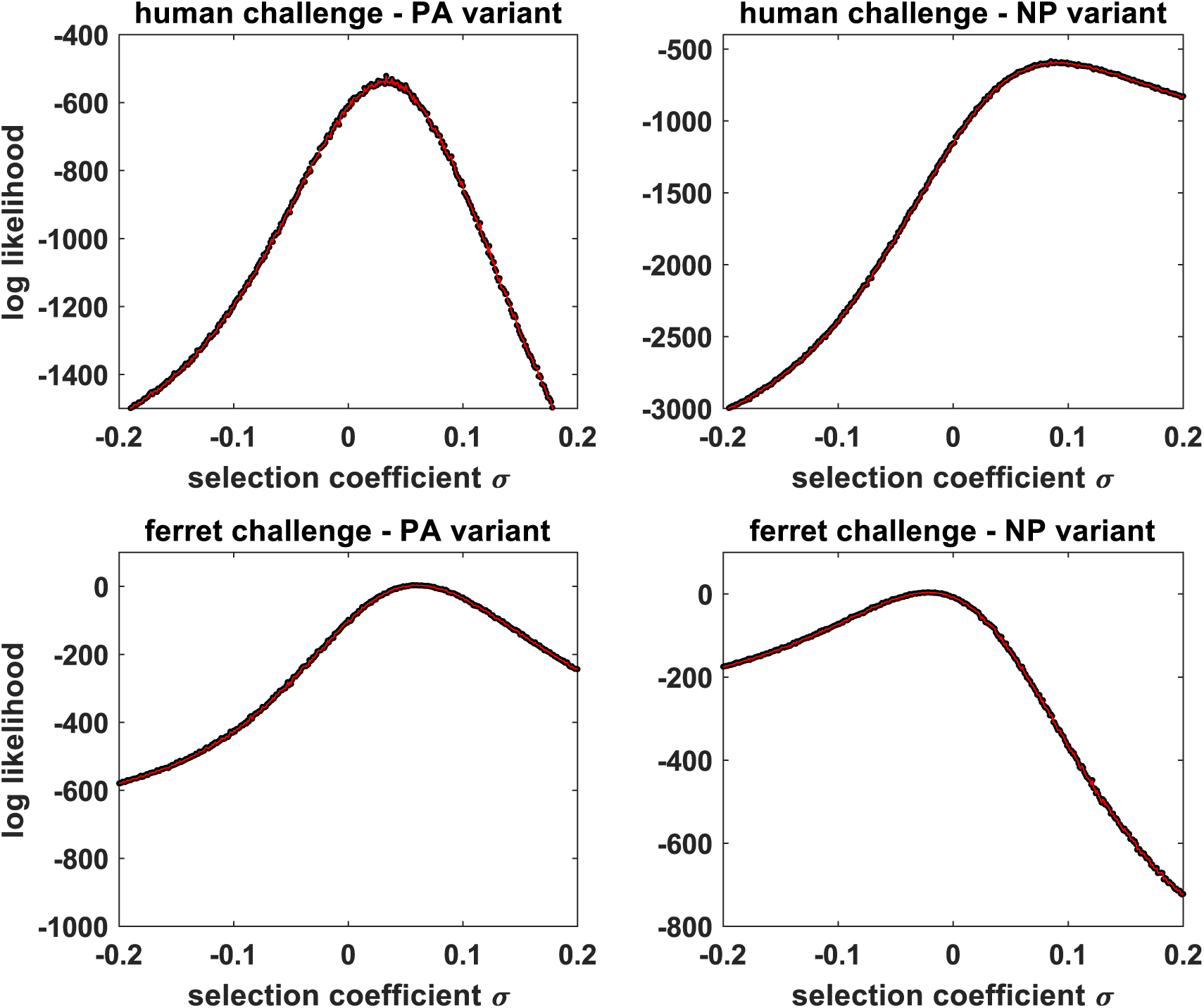
Log-likelihood profiles for selection coefficients, by host and variant for the selection-only model. (A) Estimates based on frequency changes of PA 211M-228N observed in the 13 participants. (B) Estimates based on frequency changes of NP 101D-136I observed in the 13 participants. (C) Estimates based on frequency changes of PA 211M-228N observed in directly inoculated (donor) ferrets. (D) Estimates based on frequency changes of NP 101D-136I observed in directly inoculated (donor) ferrets.

**Supplementary table 1.** Minor Variant detection in inoculum. Column Segment shows influenza virus segment as 1 = PB2, 2 = PB1, 3 = PA, 4 = HA, 5 = NP, 6 = NA, 7 = M, 8 = NS. Position denotes nucleotide position within the segment relative to reference sequence. Nucleotide_ref is the nucleotide in the reference while Nucleotide_alt is the alternative variant detected in the analysis. The column SYN shows values of “Y” when codon is maintained and “N” when mutation leads to amino acid change. The column AminoAcid_ref and AminoAcid_alt show the impact of the nucleotide mutations at the amino acid level. The column Frequency and Depth show the proportion of the variant relative to the reference and the total number of reads for the position respectively. The column Sample refers to the sample ID in the sequencing run.

**Supplementary table 2.** Variant frequency per participant during infection. Rep_num refers to the sequencing replicate. Segment_Position_AA refers to the amino acid change for that particular nucleotide position in the given segment (3 = PA and 5 = NP). Depth refers to sequencing depth. The NP value in Sample_type refers to nasopharynx.

**Supplementary table 3.** Variant frequency in ferrets during infection. Segment_Position_AA refers to the amino acid change for that particular nucleotide position in the given segment (3 = PA and 5 = NP). Depth refers to sequencing depth. The NP value in Sample_type refers to nasopharynx.

**Supplementary table 4.** Statistical analysis of evolutionary models.

| Challenge study | Variant | Model | $N_E$ | $\sigma$ | max logL | $k$ | $n$ | $AIC_c$ | $\Delta AIC_c$ |
| --- | --- | --- | --- | --- | --- | --- | --- | --- | --- |
| Human | PA 211M-228N | Selection and genetic drift | 19 | 0.070 | 13.16 | 2 | 32 | -21.90 | 0.00 |
|  |  | Selection only | Inf | 0.033 | -521.20 | 1 | 32 | 1044.53 | 1066.43 |
|  |  | Genetic drift only | 17 | 0 | 11.13 | 1 | 32 | -20.13 | 1.77 |
| Human | NP 101D-136I | Selection and genetic drift | 15 | 0.190 | 16.95 | 2 | 32 | -29.48 | 0.00 |
|  |  | Selection only | Inf | 0.085 | -584.43 | 1 | 32 | 1170.99 | 1200.47 |
|  |  | Genetic drift only | 12 | 0 | 9.64 | 1 | 32 | -17.14 | 12.35 |
| Ferret | PA 211M-228N | Selection and genetic drift | 168 | 0.065 | 8.49 | 2 | 12 | -11.64 | 0.00 |
|  |  | Selection only | Inf | 0.056 | 3.94 | 1 | 12 | -5.47 | 6.17 |
|  |  | Genetic drift only | 68 | 0 | 4.15 | 1 | 12 | -5.90 | 5.74 |
| Ferret | NP 101D-136I | Selection and genetic drift | 158 | -0.035 | 8.96 | 2 | 12 | -12.59 | 0.00 |
|  |  | Selection only | Inf | -0.022 | 4.14 | 1 | 12 | -5.88 | 6.71 |
|  |  | Genetic drift only | 127 | 0 | 7.46 | 1 | 12 | -12.52 | 0.07 |

The Akaike Information Criterion corrected for small sample size ( $AIC_c$ ) was used to
statistically compare models. The most preferred model is the one with the lowest $AIC_c$ .
This was achieved by the model including both selection and drift, for both variants and for
both hosts. The selection-only model was parameterized with a value of $N_E$ that was set to
100,000 (approximating Infinity). The genetic drift-only model was parameterized with a
value of $\sigma$ that was set to 0.

## Notes

### Competing Interest Statement

The authors have declared no competing interest.

## References

Amato, K. A., Haddock, L. A., 3rd, Braun, K. M., Meliopoulos, V., Livingston, B., Honce, R., … Mehle, A. (2022). Influenza A virus undergoes compartmentalized replication in vivo dominated by stochastic bottlenecks. Nat Commun, 13(1), 3416. doi:10.1038/s41467-022-31147-0

Bacsik, D. J., Dadonaite, B., Butler, A., Greaney, A. J., Heaton, N. S., & Bloom, J. D. (2023). Influenza virus transcription and progeny production are poorly correlated in single cells. Elife, 12. doi:10.7554/eLife.86852

Belser, J. A., Katz, J. M., & Tumpey, T. M. (2011). The ferret as a model organism to study influenza A virus infection. Dis Model Mech, 4(5), 575–579. doi:10.1242/dmm.007823

Belser, J. A., Lau, E. H. Y., Barclay, W., Barr, I. G., Chen, H., Fouchier, R. A. M., … Working group on the standardization of the ferret model for influenza risk, a. (2022). Robustness of the Ferret Model for Influenza Risk Assessment Studies: a Cross-Laboratory Exercise. mBio, 13(4), e0117422. doi:10.1128/mbio.01174-22

Bendall, E. E., Zhu, Y., Fitzsimmons, W. J., Rolfes, M., Mellis, A., Halasa, N., … Lauring, A. S. (2024). Influenza A virus within-host evolution and positive selection in a densely sampled household cohort over three seasons. Virus Evol, 10(1), veae084. doi:10.1093/ve/veae084

Biggerstaff, M., Jhung, M. A., Reed, C., Fry, A. M., Balluz, L., & Finelli, L. (2014). Influenza-like illness, the time to seek healthcare, and influenza antiviral receipt during the 2010-2011 influenza season-United States. J Infect Dis, 210(4), 535–544. doi:10.1093/infdis/jiu224

Bouvier, N. M., & Lowen, A. C. (2010). Animal Models for Influenza Virus Pathogenesis and Transmission. Viruses, 2(8), 1530–1563. doi:10.3390/v20801530

Brooke, C. B., Ince, W. L., Wrammert, J., Ahmed, R., Wilson, P. C., Bennink, J. R., & Yewdell, J. W. (2013). Most influenza a virions fail to express at least one essential viral protein. J Virol, 87(6), 3155–3162. doi:10.1128/JVI.02284-12

Carrat, F., Vergu, E., Ferguson, N. M., Lemaitre, M., Cauchemez, S., Leach, S., & Valleron, A. J. (2008). Time lines of infection and disease in human influenza: a review of volunteer challenge studies. Am J Epidemiol, 167(7), 775–785. doi:10.1093/aje/kwm375

Chopin, N. P., O. (2020). An Introduction to Sequential Monte Carlo.

Debbink, K., McCrone, J. T., Petrie, J. G., Truscon, R., Johnson, E., Mantlo, E. K., … Lauring, A. S. (2017). Vaccination has minimal impact on the intrahost diversity of H3N2 influenza viruses. PLoS Pathog, 13(1), e1006194. doi:10.1371/journal.ppat.1006194

Dinis, J. M., Florek, K. R., Fatola, O. O., Moncla, L. H., Mutschler, J. P., Charlier, O. K., … Friedrich, T. C. (2016). Deep Sequencing Reveals Potential Antigenic Variants at Low Frequencies in Influenza A Virus-Infected Humans. J Virol, 90(7), 3355–3365. doi:10.1128/JVI.03248-15

Doran, A. G., & Creevey, C. J. (2013). Snpdat: easy and rapid annotation of results from de novo snp discovery projects for model and non-model organisms. BMC Bioinformatics, 14, 45. doi:10.1186/1471-2105-14-45

Ferreri, L. M., Seibert, B., Caceres, C. J., Patatanian, K., Holmes, K. E., Gay, L. C., … Lowen, A. C. (2025). Dispersal of influenza virus populations within the respiratory tract shapes their evolutionary potential. Proc Natl Acad Sci U S A, 122(4), e2419985122. doi:10.1073/pnas.2419985122

Ferreri, L. M., & Vargas-Maldonado, N. (2025). Within-host adaptive evolution is limited by genetic drift in experimental human influenza A virus infections. Sequence Read Archive, PRJNA1367170 PRJNA1196946.

Fitch, W. M., Leiter, J. M., Li, X. Q., & Palese, P. (1991). Positive Darwinian evolution in human influenza A viruses. Proc Natl Acad Sci U S A, 88(10), 4270–4274. doi:10.1073/pnas.88.10.4270

Fullen, D. J., Noulin, N., Catchpole, A., Fathi, H., Murray, E. J., Mann, A., … Lambkin-Williams, R. (2016). Accelerating Influenza Research: Vaccines, Antivirals, Immunomodulators and Monoclonal Antibodies. The Manufacture of a New Wild-Type H3N2 Virus for the Human Viral Challenge Model. PLoS One, 11(1), e0145902. doi:10.1371/journal.pone.0145902

Ghedin, E., Holmes, E. C., DePasse, J. V., Pinilla, L. T., Fitch, A., Hamelin, M. E., … Boivin, G. (2012). Presence of oseltamivir-resistant pandemic A/H1N1 minor variants before drug therapy with subsequent selection and transmission. J Infect Dis, 206(10), 1504–1511. doi:10.1093/infdis/jis571

Gifford, D. R., de Visser, J. A., & Wahl, L. M. (2013). Model and test in a fungus of the probability that beneficial mutations survive drift. Biol Lett, 9(1), 20120310. doi:10.1098/rsbl.2012.0310

Han, A. X., Felix Garza, Z. C., Welkers, M. R., Vigeveno, R. M., Tran, N. D., Le, T. Q. M., … Russell, C. A. (2021). Within-host evolutionary dynamics of seasonal and pandemic human influenza A viruses in young children. Elife, 10. doi:10.7554/eLife.68917

Heldt, F. S., Kupke, S. Y., Dorl, S., Reichl, U., & Frensing, T. (2015). Single-cell analysis and stochastic modelling unveil large cell-to-cell variability in influenza A virus infection. Nat Commun, 6, 8938. doi:10.1038/ncomms9938

Herfst, S., Schrauwen, E. J., Linster, M., Chutinimitkul, S., de Wit, E., Munster, V. J., … Fouchier, R. A. (2012). Airborne transmission of influenza A/H5N1 virus between ferrets. Science, 336(6088), 1534–1541. doi:10.1126/science.1213362

Imai, M., Watanabe, T., Hatta, M., Das, S. C., Ozawa, M., Shinya, K., … Kawaoka, Y. (2012). Experimental adaptation of an influenza H5 HA confers respiratory droplet transmission to a reassortant H5 HA/H1N1 virus in ferrets. Nature, 486(7403), 420–428. doi:10.1038/nature10831

Jacobs, N. T., Onuoha, N. O., Antia, A., Steel, J., Antia, R., & Lowen, A. C. (2019). Incomplete influenza A virus genomes occur frequently but are readily complemented during localized viral spread. Nat Commun, 10(1), 3526. doi:10.1038/s41467-019-11428-x

Le Sage, V., Souza, C. K., Rockey, N. C., Shephard, M., Zanella, G. C., Arruda, B., … Lakdawala, S. S. (2025). Eurasian 1C swine influenza A virus exhibits high pandemic risk traits. Emerg Microbes Infect, 14(1), 2492210. doi:10.1080/22221751.2025.2492210

Li, H., & Durbin, R. (2009). Fast and accurate short read alignment with Burrows-Wheeler transform. Bioinformatics, 25(14), 1754–1760. doi:10.1093/bioinformatics/btp324

McCrone, J. T., & Lauring, A. S. (2016). Measurements of Intrahost Viral Diversity Are Extremely Sensitive to Systematic Errors in Variant Calling. J Virol, 90(15), 6884–6895. doi:10.1128/JVI.00667-16

McCrone, J. T., Woods, R. J., Martin, E. T., Malosh, R. E., Monto, A. S., & Lauring, A. S. (2018). Stochastic processes constrain the within and between host evolution of influenza virus. Elife, 7. doi:10.7554/eLife.35962

McCrone, J. T., Woods, R. J., Monto, A. S., Martin, E. T., & Lauring, A. S. (2020). doi:10.1101/2020.10.24.353748

Moncla, L. H., Zhong, G., Nelson, C. W., Dinis, J. M., Mutschler, J., Hughes, A. L., … Friedrich, T. C. (2016). Selective Bottlenecks Shape Evolutionary Pathways Taken during Mammalian Adaptation of a 1918-like Avian Influenza Virus. Cell Host Microbe, 19(2), 169–180. doi:10.1016/j.chom.2016.01.011

Pybus, O. G., & Rambaut, A. (2009). Evolutionary analysis of the dynamics of viral infectious disease. Nat Rev Genet, 10(8), 540–550. doi:10.1038/nrg2583

Rambaut, A., Pybus, O. G., Nelson, M. I., Viboud, C., Taubenberger, J. K., & Holmes, E. C. (2008). The genomic and epidemiological dynamics of human influenza A virus. Nature, 453(7195), 615–619. doi:10.1038/nature06945

Rogers, M. B., Song, T., Sebra, R., Greenbaum, B. D., Hamelin, M. E., Fitch, A., … Ghedin, E. (2015). Intrahost dynamics of antiviral resistance in influenza A virus reflect complex patterns of segment linkage, reassortment, and natural selection. mBio, 6(2). doi:10.1128/mBio.02464-14

Russell, A. B., Elshina, E., Kowalsky, J. R., Te Velthuis, A. J. W., & Bloom, J. D. (2019). Single-Cell Virus Sequencing of Influenza Infections That Trigger Innate Immunity. J Virol, 93(14). doi:10.1128/JVI.00500-19

Shetty, N., Shephard, M. J., Rockey, N. C., Macenczak, H., Traenkner, J., Danzy, S., … Lakdawala, S. S. (2024). Influenza virus infection and aerosol shedding kinetics in a controlled human infection model. J Virol, 98(12), e0161224. doi:10.1128/jvi.01612-24

Shi, T., Harris, J. D., Martin, M. A., & Koelle, K. (2023). Transmission bottleneck size estimation from de novo viral genetic variation. bioRxiv. doi:10.1101/2023.08.14.553219

Shi, Y. T., Martin, M. A., Weissman, D. B., & Koelle, K. (2025). Genetic drift acts strongly on within-host influenza virus populations during acute infection but does not act alone. bioRxiv. doi:10.1101/2025.08.27.672713

Sims, A., Tornaletti, L. B., Jasim, S., Pirillo, C., Devlin, R., Hirst, J. C., … Hutchinson, E. (2023). Superinfection exclusion creates spatially distinct influenza virus populations. PLoS Biol, 21(2), e3001941. doi:10.1371/journal.pbio.3001941

Smith, D. J., Lapedes, A. S., de Jong, J. C., Bestebroer, T. M., Rimmelzwaan, G. F., Osterhaus, A. D., & Fouchier, R. A. (2004). Mapping the antigenic and genetic evolution of influenza virus. Science, 305(5682), 371–376. doi:10.1126/science.1097211

Sobel Leonard, A., McClain, M. T., Smith, G. J., Wentworth, D. E., Halpin, R. A., Lin, X., … Koelle, K. (2016). Deep Sequencing of Influenza A Virus from a Human Challenge Study Reveals a Selective Bottleneck and Only Limited Intrahost Genetic Diversification. J Virol, 90(24), 11247–11258. doi:10.1128/JVI.01657-16

Sun, J., Vera, J. C., Drnevich, J., Lin, Y. T., Ke, R., & Brooke, C. B. (2020). Single cell heterogeneity in influenza A virus gene expression shapes the innate antiviral response to infection. PLoS Pathog, 16(7), e1008671. doi:10.1371/journal.ppat.1008671

Thompson, A. J., Wu, N. C., Canales, A., Kikuchi, C., Zhu, X., de Toro, B. F., … Paulson, J. C. (2024). Evolution of human H3N2 influenza virus receptor specificity has substantially expanded the receptor-binding domain site. Cell Host Microbe, 32(2), 261–275 e264. doi:10.1016/j.chom.2024.01.003

Valesano, A. L., Fitzsimmons, W. J., McCrone, J. T., Petrie, J. G., Monto, A. S., Martin, E. T., & Lauring, A. S. (2020). Influenza B Viruses Exhibit Lower Within-Host Diversity than Influenza A Viruses in Human Hosts. J Virol, 94(5). doi:10.1128/JVI.01710-19

Vargas-Maldonado, N., & Ferreri, L. M. (2025). Within-host adaptive evolution is limited by genetic drift in experimental human influenza A virus infections. FigShare. doi:10.6084/m9.figshare.30801677

Vargas-Maldonado, N., Shetty, N., Ferreri, L. M., Pauly, M. D., Patatanian, K., Danzy, S., … Lakdawala, S. S. (2025). Controlled human influenza infection reveals heterogeneous expulsion of infectious virus into air. medRxiv. doi:10.1101/2025.11.03.25339190

Visher, E., Whitefield, S. E., McCrone, J. T., Fitzsimmons, W., & Lauring, A. S. (2016). The Mutational Robustness of Influenza A Virus. PLoS Pathog, 12(8), e1005856. doi:10.1371/journal.ppat.1005856

Wilker, P. R., Dinis, J. M., Starrett, G., Imai, M., Hatta, M., Nelson, C. W., … Friedrich, T. C. (2013). Selection on haemagglutinin imposes a bottleneck during mammalian transmission of reassortant H5N1 influenza viruses. Nat Commun, 4, 2636. doi:10.1038/ncomms3636

Wilm, A., Aw, P. P., Bertrand, D., Yeo, G. H., Ong, S. H., Wong, C. H., … Nagarajan, N. (2012). LoFreq: a sequence-quality aware, ultra-sensitive variant caller for uncovering cell-population heterogeneity from high-throughput sequencing datasets. Nucleic Acids Res, 40(22), 11189–11201. doi:10.1093/nar/gks918

Xue, K. S., Stevens-Ayers, T., Campbell, A. P., Englund, J. A., Pergam, S. A., Boeckh, M., & Bloom, J. D. (2017). Parallel evolution of influenza across multiple spatiotemporal scales. Elife, 6. doi:10.7554/eLife.26875

